# Spatial transcriptomics reveals recasting of signalling networks in the small intestine following tissue invasion by the helminth parasite *Heligmosomoides polygyrus*

**DOI:** 10.1101/2024.02.09.579622

**Authors:** Marta Campillo Poveda, Olympia Hardy, Ross F Laidlaw, Thomas D Otto, Rick M Maizels

## Abstract

The infective larvae of the helminth *Heligmosomoides polygyrus bakeri* migrate to the small intestine, invade the submucosa, and trigger granuloma formation around each parasite. Here, we employ spatial transcriptomics to elucidate the transcriptional intricacies and cell interactions in *H. polygyrus-*infected mice. We find a generalised reduction in expression of homeostatic genes such as *Epcam*, *Pls1* (fimbrin) and *Zg16*, while cell adhesion (eg *Cldn3*, *Cdh17*) and immune-protective (*Pla2g4c*) loci are upregulated. Specific genes and cell types are associated with different spatial niches (lower crypt, upper crypt, villi and granuloma). Within the crypts, pathway analysis indicates activation of the osteopontin (*Spp1*) and pleiotrophin (*Ptn*) pathways that are poorly represented in steady-state tissues, whilst *Wnt* signalling within the crypts is abrogated by day 7 of infection. Granulomas contain concentrations of myeloid cells, NK and dendritic cells, with high expression levels of genes linked to M2 macrophages (*Arg1*, *Retlna*, *Fcer1g*) and wound repair pathways (*Reg3b* and *Mxra7*) as well as elevated *Tmbx4* that has not previously been noted. Analysis of potential ligand-receptor pairs confirmed a major complementarity between granuloma-localised SPP1 and CD44 receptors in both crypt and granuloma, as well as TGF-β/receptor interactions. Infected tissues also revealed abundant chemokine representation; among the latter category CCL6, CCL8 and MIF (macrophage migration inhibitory factor) dominated potential interactions. These results both enhance our understanding of the murine small intestine’s transcriptional landscape and also identify a new set of molecular interactions underpinning tissue-specific responses to infection that can be targeted for therapeutic intervention.

## 1. Introduction

Helminths, or parasitic worms, are amongst the most prevalent infectious agents afflicting individuals in developing nations, contributing to a global disease burden as severe as more recognized conditions like malaria and tuberculosis. Most helminth infections establish long-lived chronic parasitism, exacerbating the global health problem, and causing extensive morbidity in both humans ^1^ and livestock ^2^. A widely used model for intestinal helminth infections is *Heligmosomoides polygyrus bakeri*, a natural parasite of mice that can persist for many weeks or months in laboratory strains, defying the host’s attempts to mount an effective immune response ^3, 4^.

Intestinal helminth infections present especial challenges when parasites, such as *H. polygyrus*, invade the submucosal tissues; here the concerted efforts of macrophages and granulocytes play a pivotal role in defence against invading parasites ^5–10^. Upon oral ingestion, *H. polygyrus* larvae swiftly traverse the small intestine’s epithelial barrier, establishing in the submucosal tissue before maturing into adults and returning to the intestinal lumen 7-8 days later. Infection initiates a dominant type 2 immune response ^6, 11^, alongside an expansion of regulatory T cells ^12–14^ and early IFN-γ production ^15, 16^. Together these orchestrate crucial physiological processes, ranging from parasite containment ^17^, epithelial differentiation ^15, 18^, and restoration of barrier integrity ^19, 20^. The extensive localised and systemic properties of these responses highlight the recruitment of both “professional immune” and nonhaematopoietic cells into the type 2 effector orbit ^21^, with the larvae rapidly encased in a granuloma surrounded and infiltrated by macrophages, neutrophils, and eosinophils, among other cell types ^5, 22^. In contrast to type 1 intestinal granulomas elicited by bacterial infection through TNF and IL-1 ^23^, *H. polygyrus* granulomas are driven by type 2 cytokines and are more extensive in IL-1R-deficient mice ^24^. A type 2 effector network forms in the infected tissue, with mast cells contributing to type 2 innate lymphoid cell (ILC2) activation ^25^, upstream of Th2 differentiation ^26^, and eosinophils stimulating neuroprotective macrophages ^27^ while type 2 cytokines drive smooth muscle hyperreactivity and increased epithelial permeability ^10^.

Despite the host’s robust immune response, *H. polygyrus* establishes long-term chronic infections attributed to its immunomodulatory effects. Most notable is its secretion of proteins that mimic the function of TGF-β ^28, 29^, a pivotal regulator of the immune system through the induction of T regulatory cells, which inhibit the inflammatory effects of a variety of immune cells ^30, 31^. Until recently, investigations into helminth immuno-modulation predominantly focused on its downstream impacts on immune cell populations. However, increasing attention is being paid to the complex interactions between intestinal helminths and the epithelium ^15, 18, 32^. Particularly noteworthy are findings from the early stages of *H. polygyrus* infection, in which localised cysts or granulomas, typically 500-900 µm in size, form around each individual larva. During this phase, stem cells in the surrounding crypts have been observed to undergo a “reversal” to a foetal-like repair phenotype ^15^. Additionally, these stem cells exhibit a compromised capacity to differentiate into various effector secretory cell subsets, including tuft, goblet, and Paneth cells ^15, 18, 32^, a finding that can be recapitulated in organoid cultures of intestinal stem cells exposed to *H. polygyrus* or its secreted products ^18^.

While the interactions and effects of the nematode on the immune system and epithelium are clearly extensive, it is important to better understand the spatial context in which host-parasite interactions occur and to identify changes occurring across infected tissues that may determine parasite establishment or clearance. In recent years, a powerful new set of techniques that combine transcriptomics with histology has emerged under the “Spatial Transcriptomics” umbrella. This technology assesses gene transcription in individual cells or tissues using spatial resolution of <100 µm, accompanied by high-dimensional data collection. The visualisation and quantitative analysis of gene expression in tissue sections offers multiple insights into cell-to-cell interactions, tissue heterogeneity, and functional organisation ^33–35^, making it an impressively powerful tool to analyse the mechanistic basis of host-parasite interactions.

In this study, spatial transcriptomics were employed to investigate the localised transcriptional profile within the intestinal epithelium and lamina propria of both naïve mice and mice infected with *H. polygyrus* over the first 7 days of infection. This has allowed us to pinpoint immune cell types within the granuloma, and identify potential ligand-receptor pairs mediating communication between tissue sites in granuloma formation and stem cell differentiation. Notably, our investigation unveiled a suite of transcriptional and morphological changes extending beyond conventional immune responses, providing new insights into the complex interplay between *H. polygyrus* and the intestinal environment and how the intestinal tissue is reorganised in response to parasite invasion.

## 2. Results

### 2.1. Spatial transcriptomics confirms distinct steady state transcriptional signatures associated with different regions of the murine small intestine

When infective L3 larvae of *H. polygyrus* are ingested and migrate to the small intestine, they cross the epithelial barrier and take up residence in the submucosa for 8 days before returning to the lumen ^4^. To profile the transcriptomic landscape of the small intestine tissue, we employed the Visium (10X Genomics) platform to conduct spatial transcriptomics on formalin-fixed gut tissue, sectioned from a Swiss roll preparation (Fig. 1 A) from *H. polygyrus*-infected mice at days 3, 5 and 7 post-infection. We focused on the two fundamental categories of the small intestine, the crypt zone (including the lamina propria and the crypts) and the villi; an additional category in *H. polygyrus*-infected mice are the inflammatory granulomas forming around individual larvae within the submucosa (Fig. 1 B) which form immediately adjacent to the crypts (Fig. 1 C) and as previously reported ^5^ are rich in Type-2 macrophages expressing Arginase-1 (Fig. 1 D).

**Figure 1.**
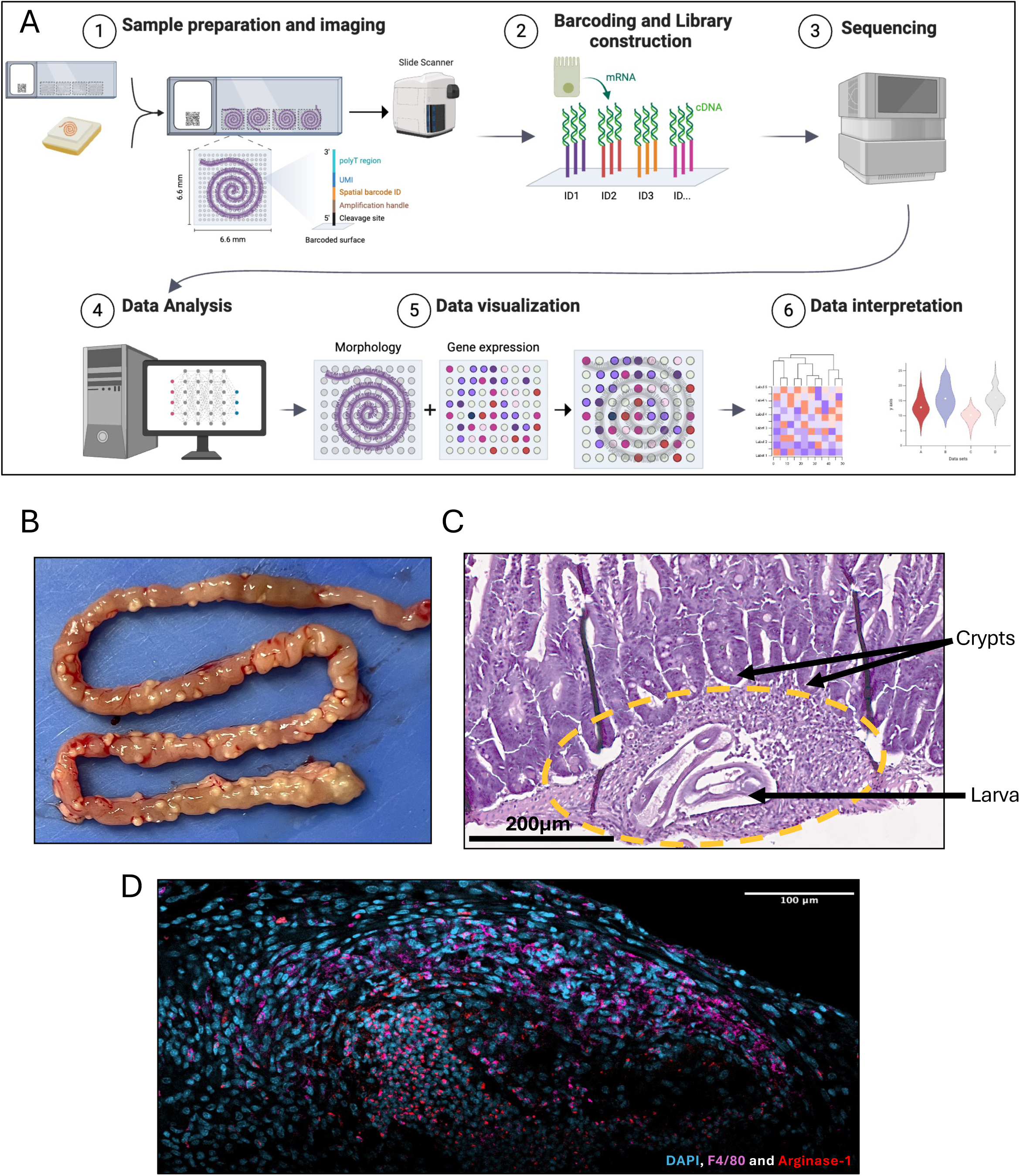
Tissue preparation and sequencing. A. Schematics of the experiments: mouse small intestine was processed for gut rolls in Formalin-Fixed Paraffin-Embedded (FFPE) blocks and further spatial transcriptomics with Visium 10X technology. B. Granulomas in the submucosal tissues of *H. polygyrus*-infected C57BL/6 female mice at day 14 following secondary infection. C. Haematoxylin and eosin (H&E)-stained section of typical granuloma containing an *H. polygyrus* larva following primary infection. D. Immunofluorescent staining of granuloma at day 7. Arginase-1-expressing macrophages in the centre of a granuloma, stained with DAPI (cyan), F4/80 (violet) and Arginase-1 (red).

We first examined naïve intestinal tissue, analysing a total of 4992 spots which were analysed and manually assigned a category, using the Loupe Browser (10X Genomics). The dataset was filtered by removing spots that were not unambiguously labelled and removing genes that had expression in 3 or fewer spots. This processed dataset from naïve mice contained expression values from 3560 valid spots across 19465 genes which had, on average, 32087 unique molecular identifiers (UMI) per spot.

The presence of known cell-specific transcriptomic markers was found to be consistent with the assigned spatial categories, with distinct sets of crypt-associated and villus-associated gene expression (Supp. Fig. 1 A). Genes such as the intestinal stem cell marker *Smoc2* ^36^ (Supp. Fig. 1 B) and growth regulator *Rack1* ^37^ define the crypt zone, where the stem cells differentiate, and developing epithelial cells start to express markers of specialisation. On the other hand, the hallmark villus signature is the villin gene *Vil1* ^38^ (Supp. Fig. 1 C). Other genes preferentially expressed by villi include the cytoskeletal components *Ezr* (Ezrin/Villin-2)*, Myo15b* (myosin XVB) and *Pls1 (Pastin1/*fimbrin), and class II major histocompatibility complex (MHC II) genes *H2-Q1* and *H2-Q2*, which help maintain the tight junctions of the enterocytes and facilitate the presentation of antigens respectively ^39^.

### 2.2 Spatial transcriptomics reveals early shift in Wnt signalling following H. polygyrus infection

We next investigated temporal shifts in signalling pathways, particularly within the crypt microenvironment at days 3, 5 and 7 following *H. polygyrus* infection, revealing extensive shifts in gene expression that are differentially regulated, with up to 700 significant changes comparing two time points (Fig. 2 A) with numerous genes prominent at each day evaluated (Fig. 2 B, Suppl. Fig 2). These changes result in a similar pattern of gene expressions on days 3 and 7 compared to naïve and day 5 (Fig. 2 B). This shift matches parasite’s life cycle events: around days 3 and 7 epithelial barrier breach and tissue penetration occurs, while Naïve and Day 5 represent the steady state and the period while the parasite is encapsulated inside the granuloma. Pathway analysis was then applied to identify key functional gene sets that are influenced during infection (Fig. 2 C). Global immune cell genes associated with CD45 became increasingly prominent over the course of infection, coinciding with the immune cell influx characteristic of early granuloma formation, where monocytes, neutrophils, and eosinophils begin surrounding the parasite ^5, 17^. Notably, increased expression of Ccl chemokines (*Ccl6, Ccl7, Ccl8*) was observed at later time points, suggesting a sustained immune recruitment process (Fig. 2 C). Given that *H. polygyrus* larvae are still developing in the submucosa at this stage, chemokine upregulation may reflect a continued attempt by the host to mount an effective immune response, while the parasite simultaneously suppresses more direct effector mechanisms ^8^.

**Figure 2.**
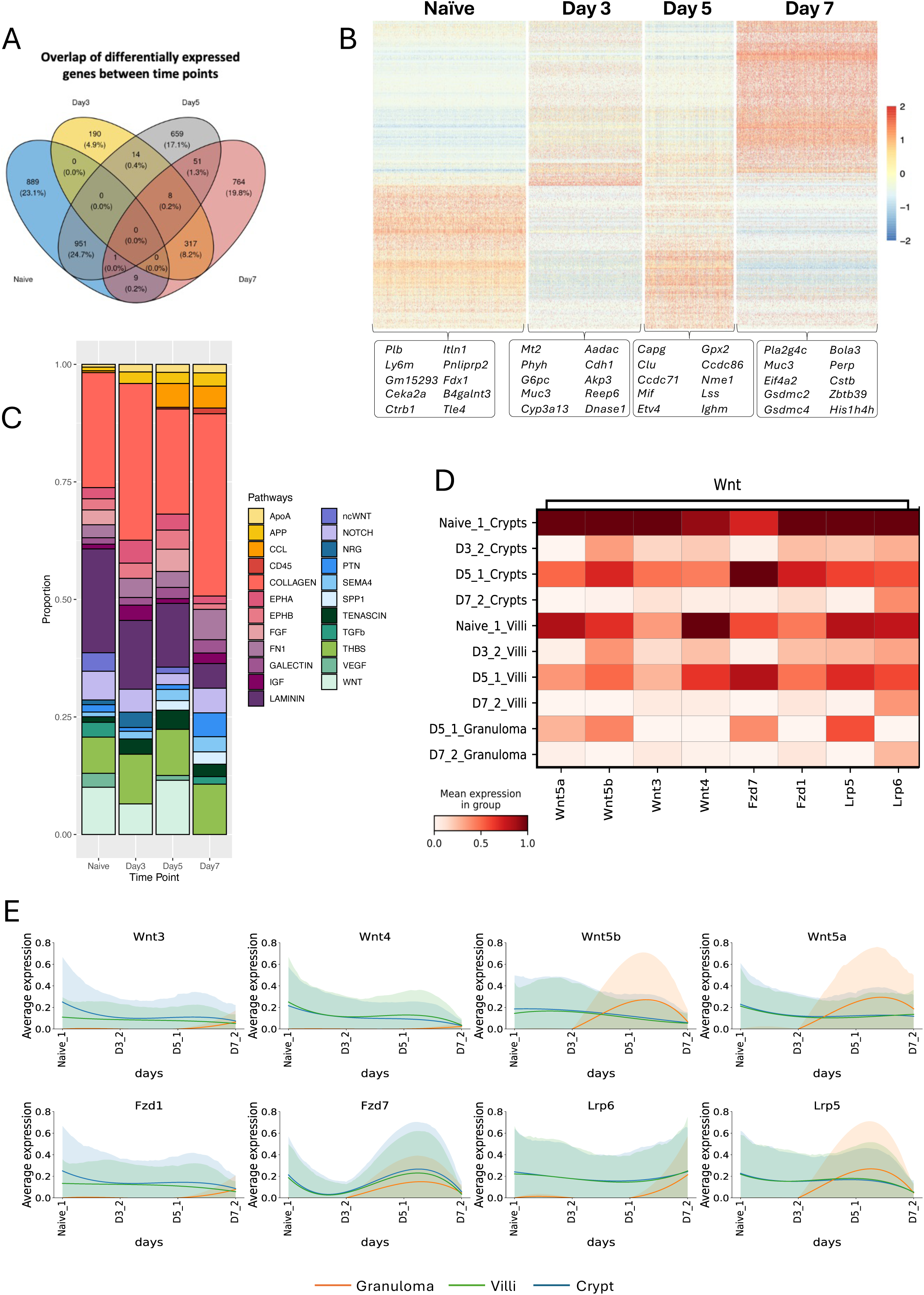
Temporal patterns of gene expression during *H. polygyrus* infection in the murine gut reveals downregulation of Wnt signalling. A. Venn diagram showing the overlap of differentially expressed genes across each timepoint. B. Heatmap of gene expression in the crypt across all timepoints. The top 10 most highly expressed genes are labelled ordered by their average log fold change values. C. Stacked barplot showing the top interaction pathways at each time point during *H. polygyrus* infection. ApoA, Apolipoproteins; APP, Amyloid Precursor Protein; CCL, Chemokines; CD45, Leukocyte Common Antigen; EPHA/EPHB, Ephrin receptors; FGF, Fibroblast Growth Factor; FN1, Fibronectin; IGF, Insulin-like Growth Factor; ncWNT, non-canonical Wnt; NRG, Neuregulin; PTN, Pleiotophin; SEAMA4, Semaphorin 4; SPP1, Secreted Phosphoprotein/Osteopontin 1; TGFb, Transforming Growth Factor-β, THBS, Thrombospondin; VEGF, Vascular Endothelial Growth Factor; WNT, Wingless/Int-1 (Integration of MMTV). D. Heatmap showing the mean expression of key ligand–receptor interactions in the Wnt signalling and TGF-β pathways across locations and timepoints. E. Average expression of each location of key ligand and receptors involved in the Wnt signalling pathway across time. Lines have been smoothed and fitted using polynomial regression, shaded regions represent confidence intervals for each fitted value. Crypts (blue), Villi (green), Granuloma (orange).

By day 7, additional signalling pathways associated with tissue remodelling became dominant, including TGF-β (*Tgfb1*), osteopontin (*Spp1*), and thrombospondins (Fig. 2 C). These factors have been implicated in fibrosis and extracellular matrix remodelling in chronic helminth infections, potentially facilitating wound healing while also reinforcing a permissive niche for parasite persistence ^28, 40^. In contrast, we noted a complete loss of Wnt pathway genes from the expression profile by day 7 post-infection (Fig. 2 C).

In naïve mice, crypt-to-crypt interactions were dominated by Wnt signalling, consistent with its role in maintaining intestinal stem cell proliferation and epithelial homeostasis ^15, 41^. This aligns with previous reports showing that in the absence of infection, Wnt-driven differentiation sustains normal epithelial renewal, balancing secretory and absorptive cell lineages ^42^. When we examined individual Wnt pathway members, a marked reduction in their expression levels was already evident by day 3 (Fig. 2 D, E). By day 7 post-infection, Wnt gene expression was almost completely absent, while Notch signalling was strongly upregulated, indicating a shift towards an absorptive lineage bias. This mirrors findings from *H. polygyrus*-infected organoid cultures, where parasite-secreted products suppress secretory cell differentiation, favouring a foetal-like repair phenotype ^18^.

### 2.3 Crypts and villi show distinct perturbations in expression profiles

We then focussed on gene expression in tissues taken 7 days post*-H. polygyrus* infection. In the infected sample, as with naïve mice described above, each spot was assigned a category by hand, using the Loupe Browser (10X Genomics), with the additional category of granuloma. Filtering of the sample was carried out in a similar fashion to the naïve, with the resulting dataset having 3964 valid spots across 19465 genes, and a mean UMI count of 10077 per spot.

When combining the overall transcriptional datasets irrespective of infection status we observed a clear division of the spots assigned to the different categories, separating the crypt zones and the villi in both naïve (Fig. 3 A) and day 7-infected (Fig. 3 B) mice, with the latter also displaying prominent granulomas. Next, when the two datasets are integrated, we found a clear separation between naïve and infected samples (Fig. 3 C). The latter finding underscores the profound transformation of gene expression in the small intestine following *H. polygyrus* infection in mice.

**Figure 3.**
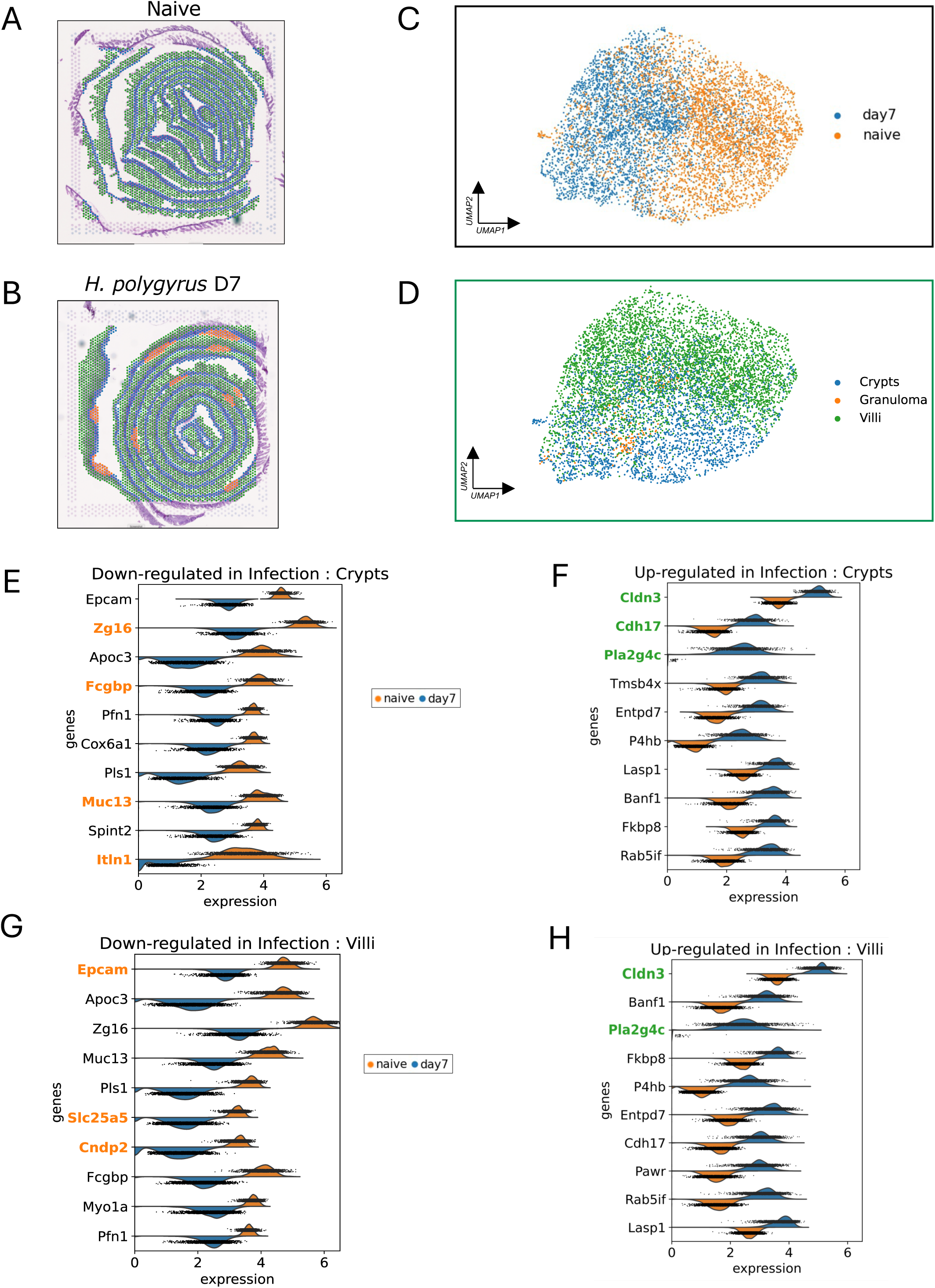
Transcriptomic differences between naïve and 7-day *H. polygyrus* infected mouse intestine. A, B Spatial plots of naïve (**A**) and day 7 *H. polygyrus* infection (**B**) highlighting tissue location clusters. C. UMAP based on Harmony integration of naïve and 7 days post *H. polygyrus* infection Visium datasets coloured by the sample origin of each of the dots. D. UMAP based on Harmony integration of naïve and 7 days post *H. polygyrus* infection Visium datasets, coloured by the tissue location of the spots. E-H. Violin plots of the expression of the top 10 differentially expressed genes in naïve and day 7 crypt (**E, F**) and villi (**G, H**). Down-regulated genes are shown in **E** and **G**, up-regulated genes in **F** and **H**, with normalized expression of naïve spots in orange, and day 7 post infection spots in blue.

We then focussed separately on the crypt and villous areas of the small intestine in naïve and infected murine tissues and identified specific genetic signatures in each of the tissue niches during infection (Fig. 3 D). Specifically, after 7 days of infection, crypts show ablated expression of genes associated with intestinal homeostasis like *Zg16*, *Muc13, Itln1* and *Fcgbp*, which help protect the intestine against bacterial colonisation ^43–45^ (Fig. 3 E, Suppl. Fig. 3 A). In the infected crypts there is also a reduction in factors that maintain the integrity of the intestinal barrier such as *Epcam* and *Pls1* ^46^.

On the other hand, a different suite of genes is upregulated in the infected crypts (Fig. 3 F, Suppl. Fig. 3 A), with most elevated expression of cell adhesion proteins (*Cldn3*, *Cdh17*) ^47, 48^ which may signify epithelial cell proliferation and modification in tight junctions at this early stage of infection. A marked induction is seen of the phospholipase A2 family member *Pla2g4c,* which is involved in and required for killing of larval *H. polygyrus* ^49^. A further induction from a low basal level is seen with the pyroptosis associated factor *Gsdmc4* (Suppl. Fig. 3 A), which is known to be upregulated in IL-4 treated and *Nippostrongylus brasiliensis* infected mouse intestinal cells ^50^.

Within the villi tissues, a similar reduction in expression profile is seen for pro-homeostatic genes such as *Epcam*, but distinct from crypt cells, the villi show down-regulation of metabolic mediators such as the mitochondrial ATP translocase *Slc25a5*, and the dipeptidase *Cndp2* (Fig. 3, Suppl. Fig. 3 B), while upregulating *Cldn3* and *Pla2g4c* as observed for crypt tissues (Fig. 3 H, Suppl. Fig. 3 B).

We also examined expression of a gene set associated with epithelial cell differentiation (*Atoh1, Gfi1, HES1* and *Neurog3*), goblet cell expression (*Clca1, Muc2*) and tuft cell function (*Alox5, Hgpds and Ptgs1*), as earlier *in vitro* studies had reported modulation in organoids treated with *H. polygyrus* products ^18^. However, while *Clca1, Hes1* and *Muc2* were reduced in infected crypts, most other gene transcripts were expressed at too low a level to be conclusive (Suppl. Fig. 3 C); a similar profile of low transcript levels apart from the same 3 genes was found In the analysis of villi (Suppl. Fig. 3 D).

### 2.4 Molecular characterization of the H. polygyrus granuloma and the surrounding small intestine

Next, we focused on the post-infection granulomas surrounding larval parasites in the submucosal tissue (Fig. 1 B-D), of which 10 distinct sites were identified (Suppl. Fig. 4 A). We compared the combined transcriptomic signatures of all granulomas in comparison to the rest of the intestinal tissue (Fig. 4 A). Remarkably, we found high expression of *Tmsb4x*, encoding thymosin beta-4, a small protein that may promote dendritic cell differentiation ^51^. As previously noted, *Arg1* (arginase-1) is highly expressed in the granulomas ^5^, as is *Retnla* (encoding RELM-α), and both genes are closely associated with alternatively activated (M2) macrophages; the marked elevation of *Ccl8* (the chemokine CCL8/monocyte chemoattractant protein-2) and *Ccl9* (CCL9/macrophage inflammatory protein 1-γ) is consistent with dominant infiltration by macrophages. Similarly, *Fcer1g,* the IgE receptor, has previously been linked to macrophages in multiple tumour and inflammatory settings, with a general activating role. Additional upregulated genes are involved in extracellular matrix (ECM) deposition (*Fn1*, *Col1a1*, and *Ctsb*) as well as antigen presentation and immune stimulation (*C1qa*, *Tnfaip2*), and lipid metabolism and oxidative stress regulation (*Apoe, Cyba, Psap*) ^52, 53^. These upregulated genes collectively imply a coordinated response involving myeloid immune cells, tissue repair, and activation of inflammatory and remodelling processes within the granuloma microenvironment.

**Figure 4.**
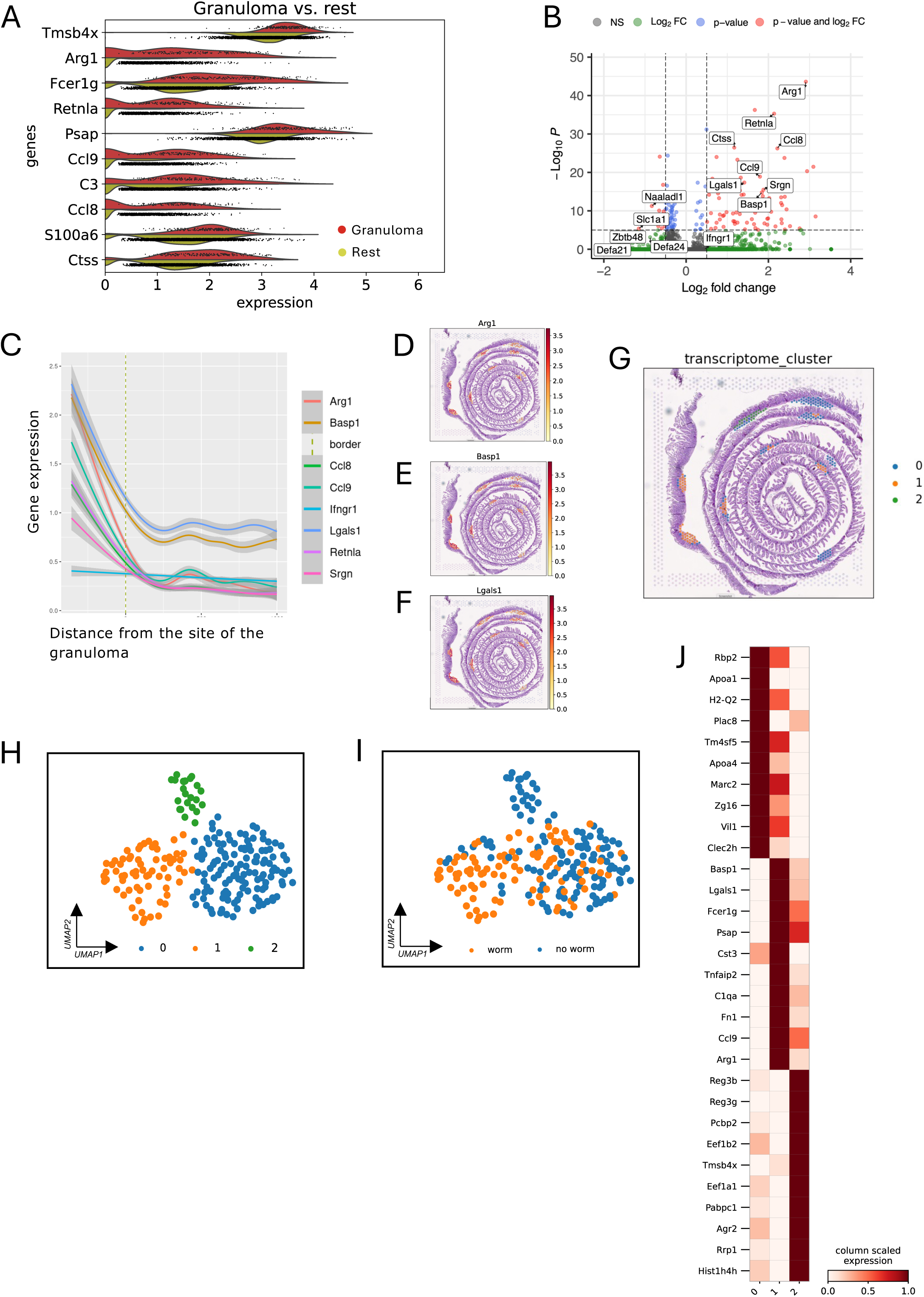
Characterizing transcriptomic signatures of granulomas in the intestine of *H. polygyrus* infected mice. A. Results of Scanpy marker gene analysis of the top 10 scoring genes for granuloma 7 days post *H. polygyrus* infection, displayed as split violin plots of normalized expression values, with red plots for granuloma spots and yellow for non-granuloma spots. B. Volcano plot showing the top 5 up/down regulated genes in the granuloma niche compared to the crypt niche. C. Spatial distribution plot showing the gene expression of the top 8 upregulated genes in the surrounding granuloma niche at d7 of infection. The dotted line denotes the boundary of the spots that are labelled as granuloma but neighbour non-granuloma spots. D-F. Spatial plots showing the gene expression of *Arg1*, *Basp1* and Lgals1 localised exclusively to the granuloma niches. G. Spatial plot of mouse intestine 7 days post infection with *H. polygyrus* with spots coloured by transcriptome-based Leiden clusters. H, I. UMAP of granuloma spots from the mouse intestine 7 days post infection with *H. polygyrus* with spots coloured by transcriptome-based Leiden clusters (H) and by absence or presence of *H. polygyrus* based on histological annotation (I). J. Scaled expression of the top 10 gene markers for each granuloma transcriptome-based Leiden cluster.

A further comparison was then made between granulomas and the crypt sites, excluding other epithelial tissues. This analysis, presented as a volcano plot in Fig. 4 B, confirms the high levels of *Arg1* and *Retnla*, as well as the monocyte chemoattractants *Ccl8* and *Basp1*. In contrast, there is concomitant *downregulation* of defensins and *Zbtb48*. The local concentration of these products was confirmed by a spatial analysis of expression relative to distance from the site of the granuloma (Fig. 4 C), and by mapping expression of transcripts for *Arg1* (Fig. 4 D), *Basp1* (Fig. 4 E) and *Lgals1* (Fig. 4 F) onto the overall landscape. We then turned to the question of heterogeneity within the set of granulomas, which differed in their presentation, and were transcriptionally variable as shown by UMAP analysis (Suppl. Fig. 4 B), which resolved into three distinct clusters (Fig. 4 G, H). Cluster 0 spots represent granulomas in which no larva is visible, either because the adult is already in the lumen or because the section is offset with respect to the worm (Fig. 4 I, Suppl. Fig 4 C). Analysis of differential gene expression by the 3 clusters (Fig. 4 J, Suppl. Fig. 4 D), reveals interesting profiles of specific gene sets. In Cluster 0, there is a higher level of immune cell products including the MHC Class II antigen *H2-Q2*, and proteins involved in interferon responses (*Ifi27l2b*), and immune regulation (*Clec2h*). Furthermore, the upregulation of *Vil1*, *Zg16*, and *Krt20* suggests the presence of epithelial cells that may contribute to the structure within the granuloma ^54^. Interestingly, one of the upregulated genes of cluster 0 granulomas is *Zg16*, which is conversely down-regulated in infected villi (Fig. 4 G, Fig. 4 J).

The observed gene expression within Cluster 1 of granulomas containing larval parasites encompasses features of both type 1 and type 2 immunity. The upregulation of pro-inflammatory genes such as M cell products *Tnfaip2* and *Ccl9* ^55^ and IgE receptor *Fcer1g* ^56^ is observed alongside genes such as *Arg1*, *Basp1*, *C1qa*, *Fn1*, *Emilin1*, and *Psap* ^57–60^ which point towards M2 macrophage activation and associated angiogenesis, tissue repair and extracellular matrix remodelling (Fig. 4 J).

Cluster 2, which represents a single granuloma which like Cluster 0 has no visible larva (Fig. 4 J and Suppl. Fig. 4), but shows a very distinctive gene profile with a lower level of macrophage activation genes, compensated by higher *Reg3b, Reg3g* and *Agr2* expression indicating resolution and regeneration of the cellular environment ^34, 61, 62^, together with *Mxra7*, encoding Matrix remodelling associated 7 protein implicated in wound-healing ^63^. Additionally, a suite of histone genes (*Hist1h4h, Hist1h1c, Hist1h3c*) ^64^, RNA processing and translation-related genes (*Eef1a1, Eef1b2, Pcbp2, Pdcd4, Rbm5, Sf3b1*), and genes associated with protein quality control suggest reinvigorated metabolic activity ^65–67^ and recovery from stress ^68, 69^ (Fig. 4 J).

### 2.5 Gene expression perturbations in tissues proximal to larval sites

In addition to defining gene expression patterns within the granulomas, we asked whether intestinal crypts adjacent to, or distant from, the sites of larval establishment showed differential transcriptional profiles. We classified sets of crypts in each category (Fig. 5 A) and identified gene sets that were up- or down-regulated in proximal sites, as candidate genes that are modulated by the presence of the helminth larvae (Fig. 5 B, C). A number of genes upregulated in the vicinity of granulomas are macrophage-associated products also found within the granulomas such as *Retnla* and *Fcer1g*, although *Arg1* was not highlighted. Of particular interest however, we noted that 2 components of the Wnt pathway, *Dact2* and *Frat2* are locally down-regulated, consistent with the overall reduction in Wnt signalling.

**Figure 5.**
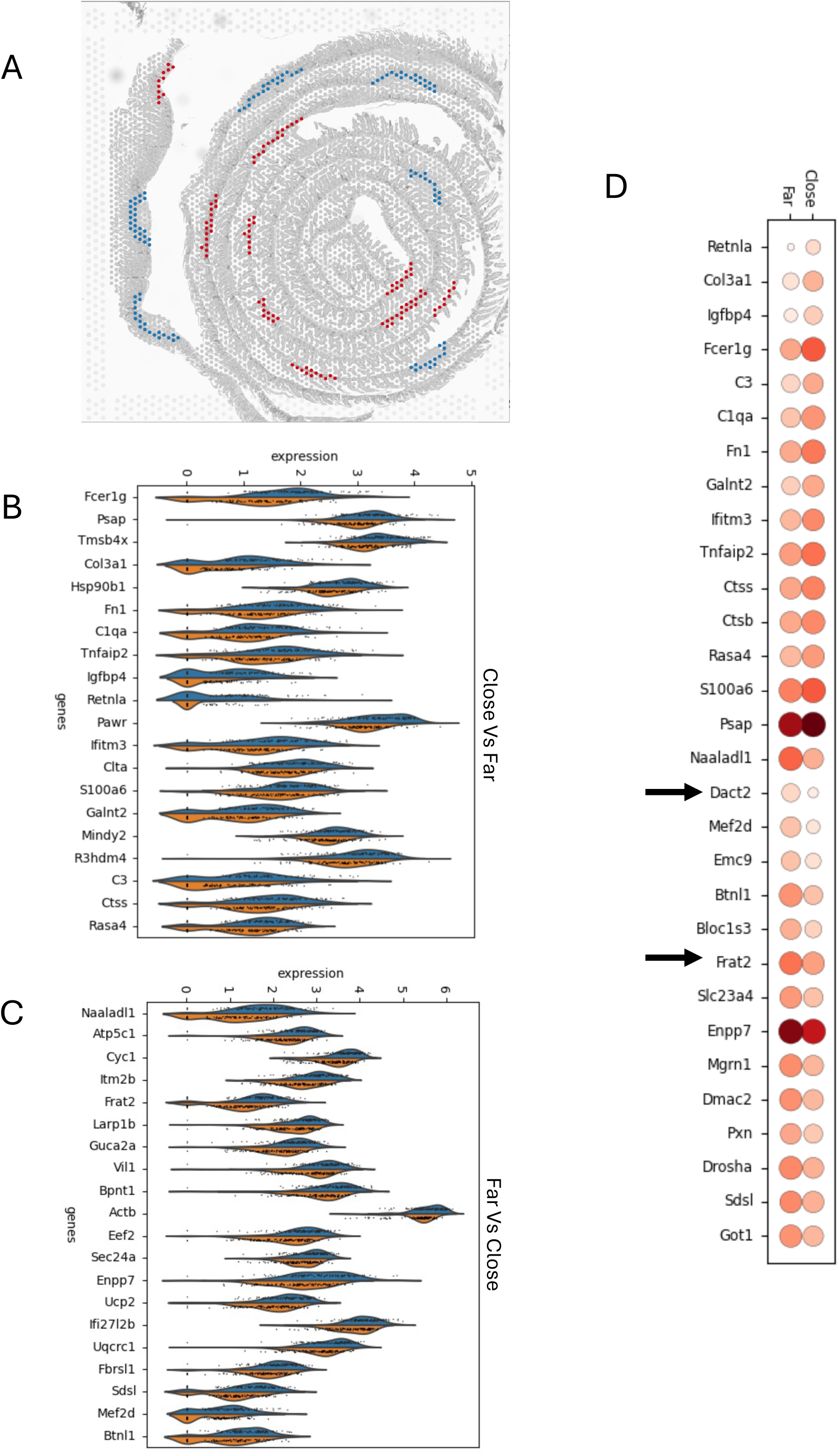
Proximity to larval parasites modulates epithelial gene expression. A. Designation of crypt areas “close” (blue) or “far (red) from sites of larval encystment. B, C. Violin plots of differential gene expression for top 20 genes upregulated (B) or downregulated (C) in crypts close to parasite locations. D. Dotplot illustrating relative gene expression levels for genes in B and C

### 2.6 Cellular composition of tissue niches

Having recreated the original biological spatial context of the intestine, we next identified the cell types present and their distribution across the spatial axis. In bulk RNA-seq data cell type abundance can be estimated by cell type deconvolution, where a scRNA-seq reference dataset is used to estimate the proportions of different cell types within a bulk sample ^70^. To deconvolute the Visium spot data, we used cell2location ^71^, a method for deconvoluting spatial transcriptomic data which builds expression profiles of cell types from a reference scRNA-seq dataset and uses these profiles to decompose the transcripts within spots into the supplied cell types from the scRNA-seq dataset.

We integrated two scRNA-seq datasets of immune and non-immune cells from the intestines of mice respectively from published studies ^72, 73^ to ensure there would be sufficient representation of the immune and epithelial cell types that comprise the intestine. Leveraging these annotations, we focused on the most proximal part of the intestine, the duodenum, which is the primary site of *H. polygyrus* tissue invasion and luminal occupancy (Fig. 6 A). Ensuring that we preserved spatial localisation of the tissue we used our calculated spatial embeddings containing our unrolled length (anterior to posterior, Fig. 6 B) and depth (lower crypt to villous tip) (Fig. 6 C) axes, and the original Visium spatial coordinates, to ensure adjacent segments in the Visium space are separated from each other (Fig. 6 D). After applying non-negative matrix factorization (see Methods) to identify which cell types colocalize together in the same spatial niches, noting that in both naïve and *H. polygyrus* infected mouse intestine, there are common signatures of cell type colocalization, especially among the non-immune cells.

**Figure 6.**
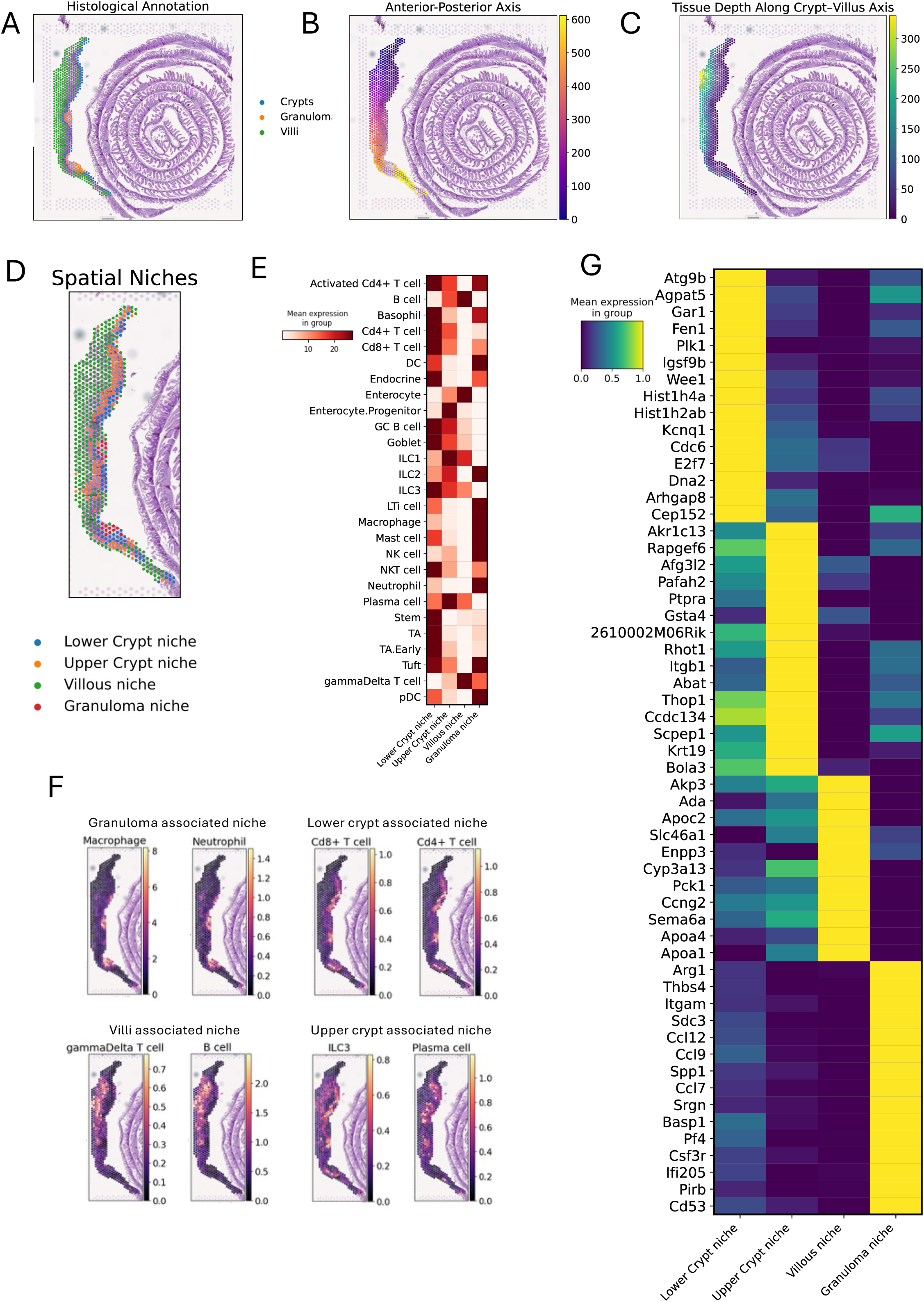
Spatial colocalization of cell types within *H. polygyrus* infected mouse intestine. A. Visium slide of mouse intestine 7 days post *H. polygyrus* coloured by the histological tissue location. B, C. Visium slides of mouse intestine 7 days post *H. polygyrus* showing spots coloured by the recreated length (B) or depth (C) axis. D. Visium slide of mouse intestine 7 days post *H. polygyrus* coloured by spatial niches. E. Heatmap showing the relative mean expression of each cell type signature present in the Xu/Huber reference single cell dataset across each spatial niche. F. Spatial projects of the top 2 predicted cell types in each spatial niche coloured by normalised cell abundance. G. Top 15 highly expressed genes for each spatial niche in the infected intestine.

Within the deepest spot layer of the crypts in both naïve and infected mice, the Lower Crypt spatial niche is dominated by transit amplifying (TA) cells and intestinal stem cells, with the presence of both CD4^+^ and CD8^+^ T cell subsets (Fig. 6 E, F, Suppl. Fig. 5 A). Directly above, the Upper Crypt is almost entirely composed of enterocyte progenitors with minor representation of lymphoid cells (Fig. 6 E, F, Suppl. Fig. 5 A). Extending toward the lumen, the villous niche colocalises enterocytes, B cells, innate lymphoid cells (ILC) 1 and 2, NK cells and γδ T cells (Fig. 6 E, F, Suppl. Fig. 5 A).

Focusing on colocalization signatures that are specific to infected mice, an accumulation of macrophage, neutrophil, plasmacytoid DC, mast cell and lymphoid tissue inducer cells localised in the intestinal granulomas (Fig. 6 E, G, Suppl. Fig. 5 A). This contrasts with the naïve condition in which macrophages and neutrophils are distributed across the intestine (Supp. Fig. 5 B,C).

While many of the cell types found to be co-localized around and within granuloma niches (macrophages, neutrophils, dendritic cells and CD4^+^ T cells) have been shown to be associated with helminth-induced granulomas, the involvement of mast cells, pDCs and Lymphoid Tissue inducer (LTi) cells has yet to be reported. Eosinophils are among the immune cell types associated with granulomas but as they are not represented in the scSeq datasets (due to high endogenous RNase expression) we were not able to distinguish them as an individual category. The presence of pDCs within the granuloma is unexpected, given their classical role as a mediator of anti-viral immune responses, but reflects highest expression of pDC markers *Siglech*, *Klk1* and *Cox6a2* ^74, 75^ in the annotated cluster (Supp. Fig. 5 D). The role of pDCs as a granuloma-associated cell type during helminth infection thus needs to be investigated.

### 2.7 Cell-to-cell interactions and signalling pathways

Following characterisation of cell types and gene expression within each spatially resolved niche in the infected murine gut, we turned to the question of interactions and molecular dialogue between these niches, by identifying complementary ligand-receptor pairings using CellChatp ^76^. From the granuloma niche, all predicted interactions were with the lower crypt, and indeed these represented the highest number of predicted interactions in the whole study (Fig. 7 A). Numerous interactions were also noted between other sites, connecting the lower crypt in particular with the upper crypt and villi that were not predicted to directly interact with granuloma products.

**Figure 7:**
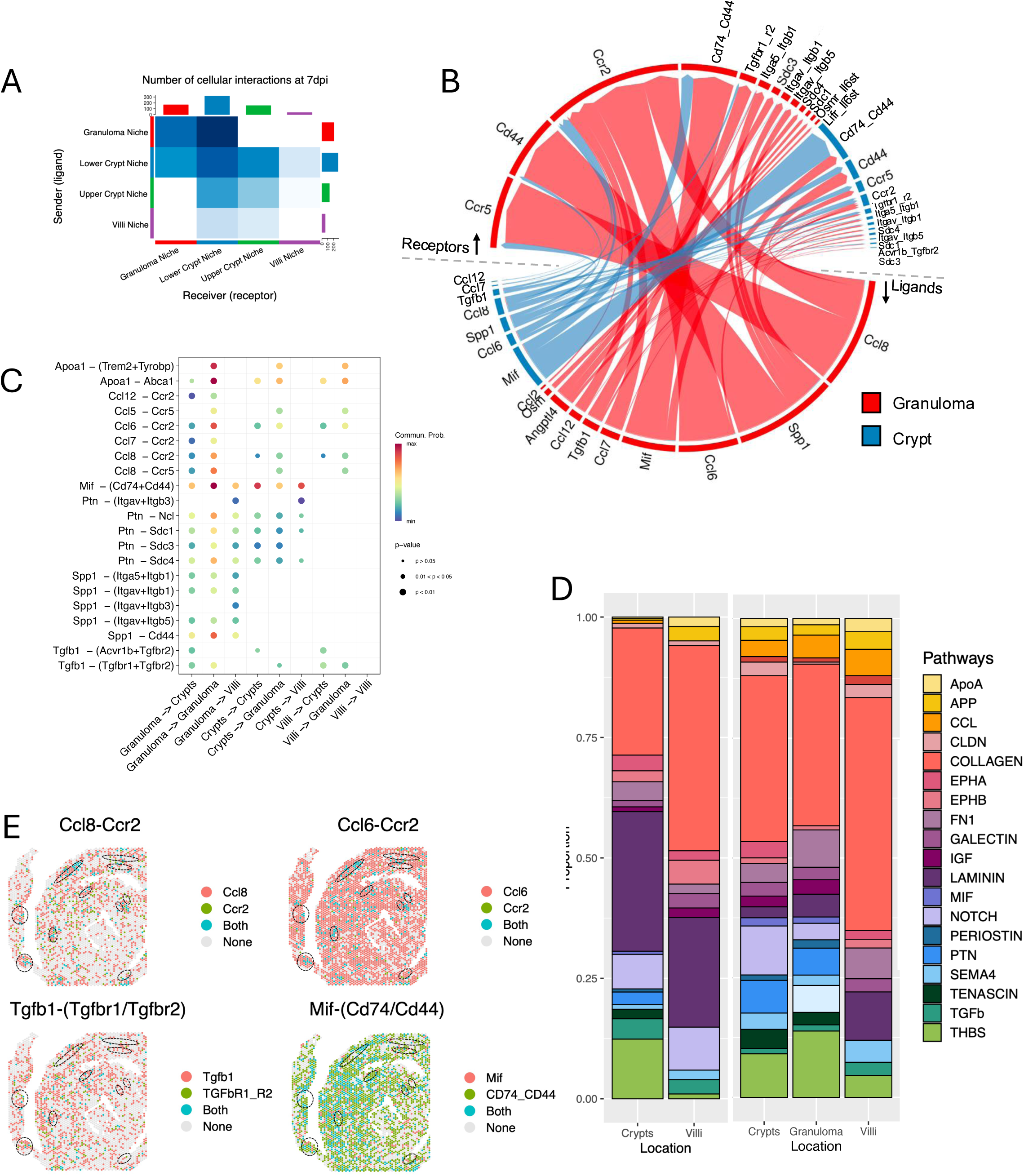
Cellular interactions across spatial niches during *H. polygyrus* infection in the murine gut. A. Heatmap showing the number of ligand-receptor (L-R) interactions inferred by CellChat for each spatial niche at 7 days post infection. Bar plots at top and side show the sum of all interactions. B. Circos plot visualisation showing upregulated ligand and receptor pairs between the granuloma (red) and the lower crypt (blue) niche. Ligands are placed in the lower half and receptors in the upper half, with arrows showing directionality of the interaction. C. Dot plot showing the communication probability of each significant ligand receptor interaction between various spatial niches in the murine gut at 7 days post infection. D. Stacked proportion bar plot showing the interaction pathways associated with each location of the murine intestine day 7 post-infection. E. Spatial projection of key interacting pairs involved between the granuloma and crypt niche during *H. polygyrus* infection, coloured by gene expression.

Enriched predicted signalling between the granuloma and lower crypt niches primarily represented immune cell activation and differentiation pathways. Dominant chemokines in the granuloma profile are CCL6, CCL7, CCL8 and CCL12, with the CCL8/CCR5 pairing with the lower crypt acting in a bidirectional manner, likely to underpin the activated myeloid and lymphoid populations recruited to the granuloma, while also able to act on chemokine receptors in the lower crypt (Fig. 7 B, C). Interestingly, the among the strongest interactions is between the chemokine-like macrophage migration inhibitory factor (MIF) expressed in both the lower crypt and the granuloma, and its receptors CD74 and CD44 present in both niches (Fig. 7 B, C), as this chemokine-like mediator is known to be essential for immunity to *H. polygyrus* as well as to the rat nematode *Nippostrongylus brasiliensis* ^77^.

As may be expected for the tissue remodelling involved in granuloma formation, there are prominent interactions with ligands for extracellular matrix and proteoglycan including pleiotrophin (*Ptn*) which binds heparin and the syndecin receptors *Sdc1, Sdc3* and *Sdc4* as well as nucleolin (*Ncl*), and the IL-6-like cytokine Oncostatin M (*Osm*) (Fig. 7 C). Particularly conspicuous is *Spp1* (secreted phosphoprotein 1, also known as sialoprotein or osteopontin) which binds CD44 and integrins such as α4 ^78^, and in a comparative pathway analysis of the 3 tissues sites, is found to be prominent in granulomas (Fig. 7 D). Spp1 can also interact with αv integrins involved in activation of latent TGF-β. Indeed, TGF-β signalling adopts a high profile in both granuloma-crypt interactions, primarily represented by TGF-β1 from the granuloma binding the canonical TGF-β receptors, but also signals from the lower crypt to the upper crypt (Fig. 7 B, C). It is known that blocking TGF-β signalling promotes expulsion of *H. polygyrus* ^79^, and most intriguingly that this parasite secretes mimics of TGF-β that bind the same receptors together with CD44 ^28, 80^. Combined mapping of key ligand-receptor pairs reveals their spatial restriction, largely to the sites of granulomas, although MIF-CD74-CD44 appears to be more generalised through the infected tissue (Fig. 7 E).

Collectively, these findings demonstrate a progressive disruption of epithelial renewal and immune-epithelial crosstalk during *H. polygyrus* infection. Early suppression of Wnt signalling (day 3) precedes Notch-dominated differentiation (day 7), a pattern that has been previously linked to helminth-induced epithelial remodelling. Concurrently, crypt-granuloma interactions evolve from inflammatory (IL-6-driven, day 5) to a tissue remodelling state (TGF-β, osteopontin, day 7), reflecting the dual nature of the host response - attempting both parasite clearance and damage repair. These results reinforce the notion that *H. polygyrus* actively reshapes its host environment, leveraging immune suppression and epithelial reprogramming to establish chronic infection. Understanding these spatially distinct and time-dependent interactions provides deeper insight into how helminths exploit the gut niche and highlight potential targets for therapeutic intervention.

## 3. Discussion

Spatial transcriptomics has emerged as a powerful tool to unravel the intricacies of gene expression patterns within the complex architecture of tissues ^81^. In this study, we employed the Visium platform to conduct spatial transcriptomics on murine small intestine tissues, providing a comprehensive view of the transcriptional landscape during steady-state conditions and in response to *H. polygyrus* infection. Furthermore, we uncover the localisation of various cell types within the context of steady state and helminth-infected intestines and infer the cell-cell interactions that take place between them.

Upon *H. polygyrus* infection, dynamic transcriptional changes were observed in the small intestine, particularly within the granuloma and surrounding areas. One such change is the apparent upregulation of tight junction genes in the helminth infected gut compared to the naive, which could impair the host’s ability to increase fluid egress as part of the ‘weep and sweep’ response against helminths. In addition, as *H. polygyrus* infection can lead to the suppression of differentiated specialist epithelial cells like goblet, Paneth and tuft cells ^15, 18, 32^, this may explain attenuated expression of specific stem cell differentiation genes of the crypt areas.

The transcriptomic analysis at day 7 post-infection revealed a multifaceted immune response, with distinct granuloma signatures categorized into clusters based on the distance to the parasite, which may be indicative of a longitudinal response to the parasite from active inflammation to resolution of the immune response. While the parasite is still present within the granuloma, a variety of type 2 immune response factors are expressed, including *Fcer1g* and products of M2 phenotype macrophages like *Arg1* and *Retlna*, all of which are associated with the clearance/expulsion of parasites. Once the parasite is no longer detected in the granuloma, there is a return towards homeostasis, with an increase in extracellular matrix remodelling and maintenance of the intestinal barrier through keratin deposition.

Despite potential limitations of deconvolution approaches, they allowed us to leverage spatially resolved neighbourhoods with existing single cell data to predict cell-to-cell interactions of spatial niches within the small intestine. Common signatures of cell type colocalization were identified, particularly in the crypts and villi, emphasizing the coordinated organization of various cell types within specific tissue regions. Specific colocalization patterns in *H. polygyrus* infection, particularly in the granulomas, highlighted the cellular interactions occurring during the host response to parasitic infection. In contrast, in the steady-state there is negligible activity from immune cells and the interactome is rather associated with epithelial cell differentiation and maintaining tissue integrity of the intestine. Thus, this landscape is dramatically altered in response to parasitic infection and the formation of granulomas.

Although our study demonstrates the heterogeneity of the host response across different spatial compartments of the intestine during *H. polygyrus* infection, the advent of higher-resolution spatial sequencing techniques would allow insight into the transcriptomic landscape of individual cells, and potentially the helminth itself, within their native spatial context. This will offer insights into cell types, their spatial distribution, potential heterogeneity within specific regions, and interactions with the parasite, enabling a comprehensive characterization of the dynamic host-pathogen interplay. Future work includes integrating additional time points of *H. polygyrus* infection in the murine intestine to map the spatial-temporal dynamics of the interplay of the host immune system and gut epithelia. These insights gained from our study lay the groundwork for future investigations into the molecular mechanisms driving host defence and tissue responses during parasitic infections. To conclude, this report highlights the main strengths of spatial transcriptomics, which is to access and unravel previously inaccessible details, paving the way for a more in-depth comprehension of the molecular and spatial dimensions of host immune responses and tissue-specific adaptations to parasitic challenges.

## 4. Materials and Methods

### 4.1 Mice and parasites

Eight-week-old female C57BL/6 mice purchased from Envigo UK and housed in individually ventilated cages were used throughout this study. All animal studies were performed under the UK Home Office Licence and approved by the University of Glasgow Animal Welfare and Ethical Review Board. Mice were infected by oral gavage with 400 infective third-stage larvae (L3) *H. polygyrus*, maintained as previously described ^82^. Mice were culled on days 3, 5 and 7 post-infection.

### 4.2 Immunohistochemistry

Gut roll sections were made using a microtome through the gut rolls at a thickness of 5 µm before mounting on glass slides. Sections were deparaffinized by immersing slides in xylene, then hydrated through 100%, 90%, and 70% ethanol successively. Heat-induced epitope retrieval was performed in citrate buffer (Thermo Fisher Scientific), and then sections were blocked using 2.5% normal horse serum blocking solution (Vector Laboratories) for 1 h at room temperature. Slides were then incubated overnight at 4°C with the corresponding Ab () in 2.5% normal horse serum blocking solution. Polyclonal rabbit IgGs (Abcam) were used as an isotype control. After washing, sections were incubated with specific AlexaFluor secondary Abs (1:1000), washed, stained with DAPI for 5 min and finally mounted using Heatshields Vectashield HardSet Antifade Mounting Medium (Vector Laboratories). Slides were imaged using a Nikon-AX inverted microscope. The resulting image files were analysed using ImageJ/Fiji.

### 4.3 Tissue processing and library preparation

Processing small intestine tissue involved the removal of the gut-associated adipose tissue following which luminal contents were removed by washing with cold PBS. Starting from the most distal end, the small intestine was mounted onto a wooden skewer with the luminal side facing outwards. After 4 h fixing in 4% NBF, tissue was cut longitudinally and rolled creating a gut roll with the distal small intestine at the centre. The roll was then left to fix overnight, followed by processing and embedding in paraffin ^83^. Five μm sections, which included at least one granuloma, were made using a microtome and placed on Visium 10x Genomics slides (PN-2000233), fitting the four 6.5mm² capture areas (Fig. 1).

For analysis, slides with small intestine tissue were stained with H&E, and imaged by NanoZoomer-SQ Digital slide scanner. Sequence libraries were processed per the manufacturer’s instructions (10x Genomics, Visium Spatial Transcriptomic). After cDNA strand synthesis, cDNA was quantified using quantitative RT-PCR ABI 7500 Fast Real-Time PCR System and analysed with ABI 7500 Software 2.3 (Fig. 1).

### 4.4 Sequencing and data processing

Visium libraries were sequenced using a P3 flow cell to a minimum of 200 million reads per sample on the NextSeq2000 instrument, paired-end 2×100bp. The sample images were then analysed and spots were assigned a metadata value according to the type of tissue the spot captured: Crypt, Villus, Peyer’s Patch and in infected tissues also Granuloma and *H. polygyrus* (referred to as ‘Worm’). Samples were mapped against the *Mus musculus* mm10 reference using 10X Genomics’ Spaceranger version 2.1.1 (10X Genomics) on default parameters except for the loupe alignment JSON file, which was edited so that unlabelled spots were also included in the final mapping output. The Spaceranger mapped Naive, D3, D5 and D7 samples underwent quality control using SCANPY ^84^, with spots with high UMI counts detected across the samples removed.

### 4.5 Comparison of crypts and villi across H. polygyrus infection

The quality controlled naïve, D3, D5 and D7 samples were concatenated together into an AnnData object, with only the genes that were detected in all four of the datasets being present in the concatenated dataset. The expression values were normalised by their total sum, and each cells normalised counts scaled to the median UMI count of the concatenated object. These values were then log1p transformed.

A series of differential expressed gene analyses were carried out across the infection time series. The concatenated object was split into two objects for spots labelled ‘Crypt’ or ‘Villi’. For these datasets, each timepoint of interest was compared to its adjacent time points. For example, the D5 sample was compared against the D3 and D7 samples, while the naive sample was compared against the D3 sample. Differentially expressed genes were defined as those with a Benjamini-Hochberg corrected p-value < 0.05, with p-values generated using a Mann Whitney U test as the default method of SCANPY.

### 4.6 Characterising granulomas within H. polygyrus-infected mice intestine

The mapped and quality controlled D5 and D7 post *H. polygyrus* infection samples were loaded into SCANPY ^84^, concatenated into a single AnnData object, and the spots designated as ‘Granuloma’ were extracted. Expression values of the spots were normalized by their total UMI counts and scaled so they added up to the median total UMI counts of the concatenated object. These values were then log1p transformed. The top 2000 variable genes were found using default parameters from SCANPY. Principal Components analysis was carried out on the normalized expression data using only the genes identified as being highly variable, and batch correction of the PCA space was carried out using Harmony. A nearest neighbours (nn) graph was constructed using the first 10 Harmony-adjusted PCs, with this graph being used as the basis for Leiden clustering (resolution = 0.4) and UMAP ^85^. Marker genes of the resulting Leiden clusters were identified using SCANPY’s rank_genes_groups function, using t-test to assess for statistical significance and Benjamini-Hochberg to adjust the p-value to correct for multiple-testing.

### 4.7 Loupe browser

Loupe Browser (v.5.0.0; 10X Genomics) is a desktop application compatible with Windows and Macintosh operating systems. It facilitates the easy visualisation and analysis of Visium gene expression data. This software is employed for the identification of significant genes and cell types, as well as for exploring substructures within cell clusters. The Loupe Browser viewer utilises single points representing cell barcodes. To visualise data within the Loupe Browser 2D space, Cell Ranger transfers Principal Components Analysis (PCA)-reduced data into a t-SNE (t-Stochastic Neighbour Embedding) plot. T-SNE is a nonlinear dimensionality reduction method with modifications by 10X Genomics. Loupe Browser includes a “Categories” mode, allowing users to label subpopulations of cells in the clustering plot with specific categories.

### 4.8 Preparing the single-cell RNA sequencing reference datasets

The raw expression matrices and metadata of the Xu *et al*. ^72^ and Haber *et al.* ^73^ scRNA-seq data were loaded into Seurat ^86^. For the Xu data, more quality control of the samples was carried out. This consisted of removing cells with remarkably high/low nFeature_counts and also those with a high percentage of mitochondrial reads per cell. The cut-off values can be seen by viewing the relevant code on our GitHub. The metadata of the Xu set were then simplified, with the cell types being designated as ‘low UMI’ being merged with their regular UMI counterparts and specific subsets of cell type e.g. ‘DC (Cd103+ Cd11b+)’, ‘DC (Cd103+ Cd11b-)’ and ‘DC (Cd103-C2)’ being simplified as just ‘DC’. Nonimmune cells were also removed from the Xu datasets. The Xu and Haber Seurat objects were then merged and converted into h5ad format.

### 4.9 Cell2location analysis of H. polygyrus infected and naïve mice intestine

For setting up the model of the Xu and Haber merged scRNA-seq dataset, we used the sequencing run of each of the datasets as the ‘batch_key’ and included the condition of the datasets (e.g. allergy, parasite infected, naive/control) as a categorical covariate. The training parameters for the training of the Visium slide model in cell2location ^86^ were chosen as follows. Within all the D3, D5, D7 and naïve Visium samples there was variation in total UMI counts that could not be explained by the tissue, thus the RNA detection sensitivity parameter was set to 20, as per the recommendation of the cell2location authors. The number of cells per location was chosen to be 50, based on visual observation of the scanned slides. The models were trained on a GPU with 80 Gb of RAM. The 5% quantile cell abundance was stored in the Visium anndata objects and used for all subsequent analysis and visualisation.

Non-negative factorization (NMF) analysis in cell2location was carried out, using the concatenation of the new length and depth coordinates and the original Visium coordinates as the spatial basis for the NMF. For the D7 sample, 7 factors were created, five factors were created for the D5 sample, while three factors were created for the naïve and D3 samples.

### 4.10 Cellular communication inference of H. polygyrus infection in the murine intestine

Spatial niches defined by the NMF factorisation analysis yielded three distinct spatially resolved neighbourhoods termed the villi, upper crypt, lower crypt and granuloma niche. Spots assigned these labels were fed into CellChat (v.2.1.1) ^76^ alongside the spatial coordinates from the full resolution tissue image to allow resulting interactions to be within spatial constraints. The conversion of spatial coordinates from pixels to micrometres was calculated using the ratio of the theoretical spot size set to 55um over the number of pixels that cover the diameter of the spot. In addition to this, the communication probability of two cells interacting was also restricted with a contact range set to 100 as recommended by the CellChat 10X Visium workflow. The CellChat database used was set to the organism ‘*mouse*’ and all functional interaction annotations were used except those classified as ‘*Non-protein Signalling*’ to avoid the inclusion of interactions involving synaptic signalling which lies outside the context of the murine intestinal tissue.

### 4.11 Package versions

Python version 3.10.12 was used with the following packages: Pandas – v.1.5.3, Numpy – v.1.23.0, SCANPY – v.1.9.3 ^76^, anndata – v.0.8.0, squidpy – v.1.3.1, scipy – v.1.10.1, sklearn – v.1.3.0, cell2location – v.0.1.3. R version v.4.2.2 was used with the following packages: Seurat – v.4.3.0, harmony – v.1.2.0, SeuratDisk – v.0.0.0.9020, CellChat – v.2.1.1. When training the cell2location models on GPUs, Python v.3.10.9 was used along with the following package versions: cell2location – v.0.1.4, scipy – v.1.10.0, SCANPY – v.1.9.6, scikit-learn – v.1.2.1, numpy – v.1.23.4, pandas – v.2.1.4.

### 4.12 Code availability

The code to replicate this analysis can be found at the following github repository: https://github.com/No2Ross/Visium_Hpolygyrus_gut

## Acknowledgements

We thank Dr Juan Quintana for expert advice and Claire Ciancia for excellent technical assistance. The authors gratefully acknowledge Glasgow Polyomics (University of Glasgow) for their support and assistance.

## Author contributions

All authors helped shape the research, analysis, and manuscript. The study was designed by M.C.P. and R.M.M. with input from T.O. M.C.P., O.H and R.F.L generated the data, performed the analysis and drafted the manuscript figures and text. R.M.M and T.O. provided critical feedback and edited the manuscript prior to submission.

