## supplemental figures for "Spatial transcriptomics reveals recasting of signalling networks in the small intestine following tissue invasion by the helminth parasite *Heligmosomoides polygyrus*"

#### Supplementary Material

**Supplementary Figure 1.**

**Characterization of steady-state mouse intestine transcriptome.**

A. Heatmap of the normalized expression values of the top 25 scoring SCANPY marker genes of crypt and villi spots from naïve mouse intestine.

B. Spatial feature plots showing the normalized expression of crypt marker *Smoc2*.

C. Spatial feature plots showing the normalized expression of villus marker *Vil1*.

**Supplementary Figure 2.**

**Temporal patterns of gene expression during the first 7 days of *H. polygyrus* infection**

A. Results of SCANPY marker gene analysis comparing naïve, d3, d5 and d7 crypt and villi displayed as heatmaps showing the normalized expression of the top 50 genes with overall upregulation over 7 days.

B. As A, but top 50 genes showing down regulation over 7 days.

**Supplementary Figure 3.**

**Comparison of crypt and villus gene expression by day 7**

A, B. Results of SCANPY marker gene analysis comparing naïve and day 7 crypt and villi displayed as heatmaps showing the normalized expression of the top 50 scoring genes in each comparison.

C, D. Analysis of genes associated with stem cell, goblet cell and tuft cell expression.

**Supplementary Figure 4.**

**H&E stain of day 7 *H. polygyrus* infected mouse intestine.**

A. Visium slide of infected mouse intestine with spots coloured by their corresponding granuloma.

B. UMAP with corresponding colours.

C. Heatmap showing normalized gene expression of the Scanpy top 25 scoring marker genes for each of the granuloma Leiden clusters.

D. Close-up images of each of the granulomas in the sample (red arrow = *H. polygyrus*).

**Supplementary Figure 5.**

**Spatial co-localisation of cell types within naive mouse intestine.**

A. Dotplot showing the contribution of different cell types to the non-negative matrix factorization factors, with each dotplot coloured by a cell types normalized cell abundance across the factors.

B. Visium slide of naïve mouse intestine coloured by the tissue metadata.

C. Visualization of distribution across tissue niche, each spot coloured by the mean UMI count for the different factors.

**Supplementary Figure 6.**

**Cell2location results for mouse duodenum at day 7 post-*H polygyrus* infection.**

A. Spatial prediction of the different immune and non-immune cell types in the mice gut presenting *H. polygyrus* granulomas. Each spot is coloured by the abundance of the cell type.

Supplementary Figure 1

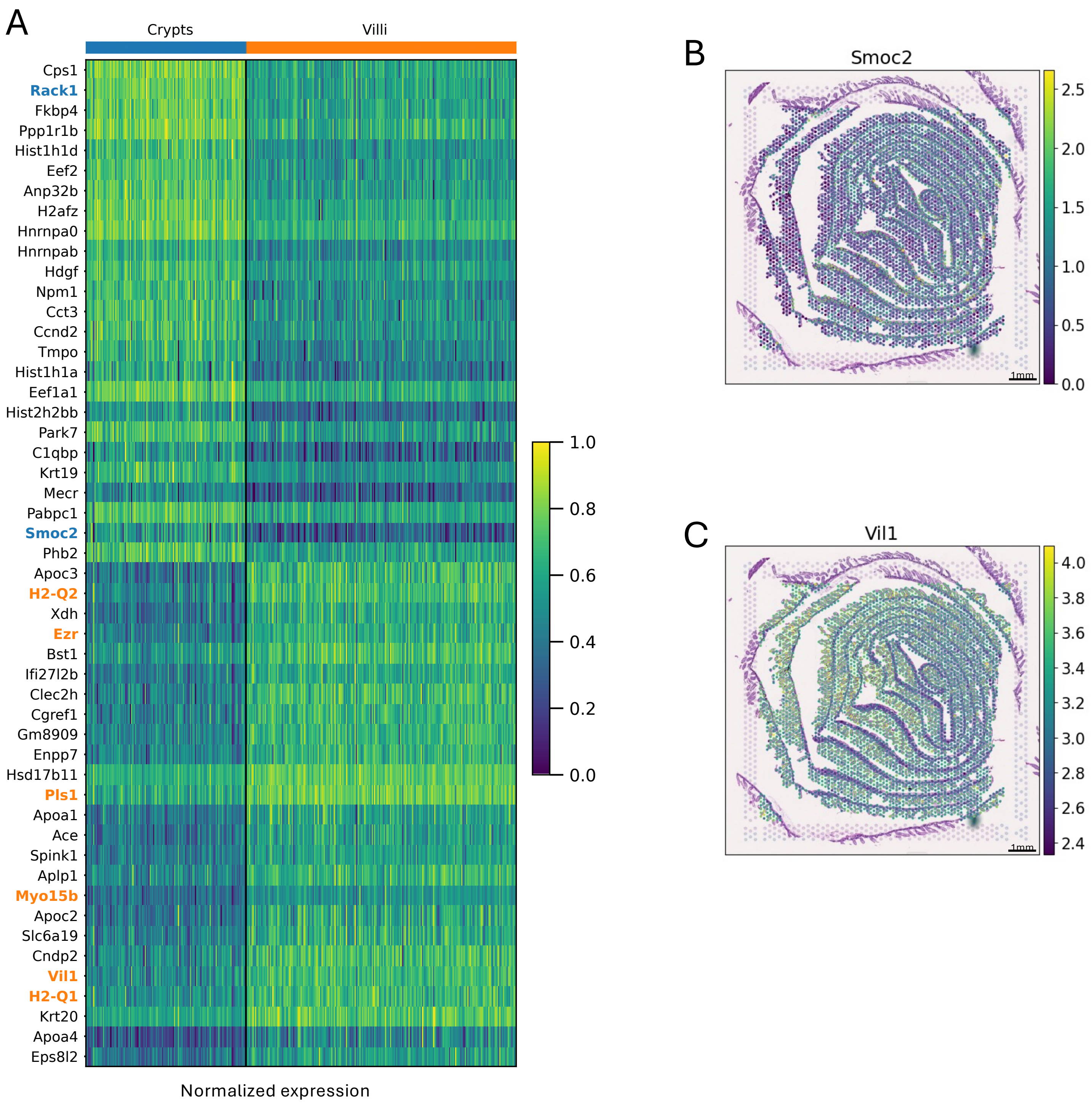

Supplementary Figure 2

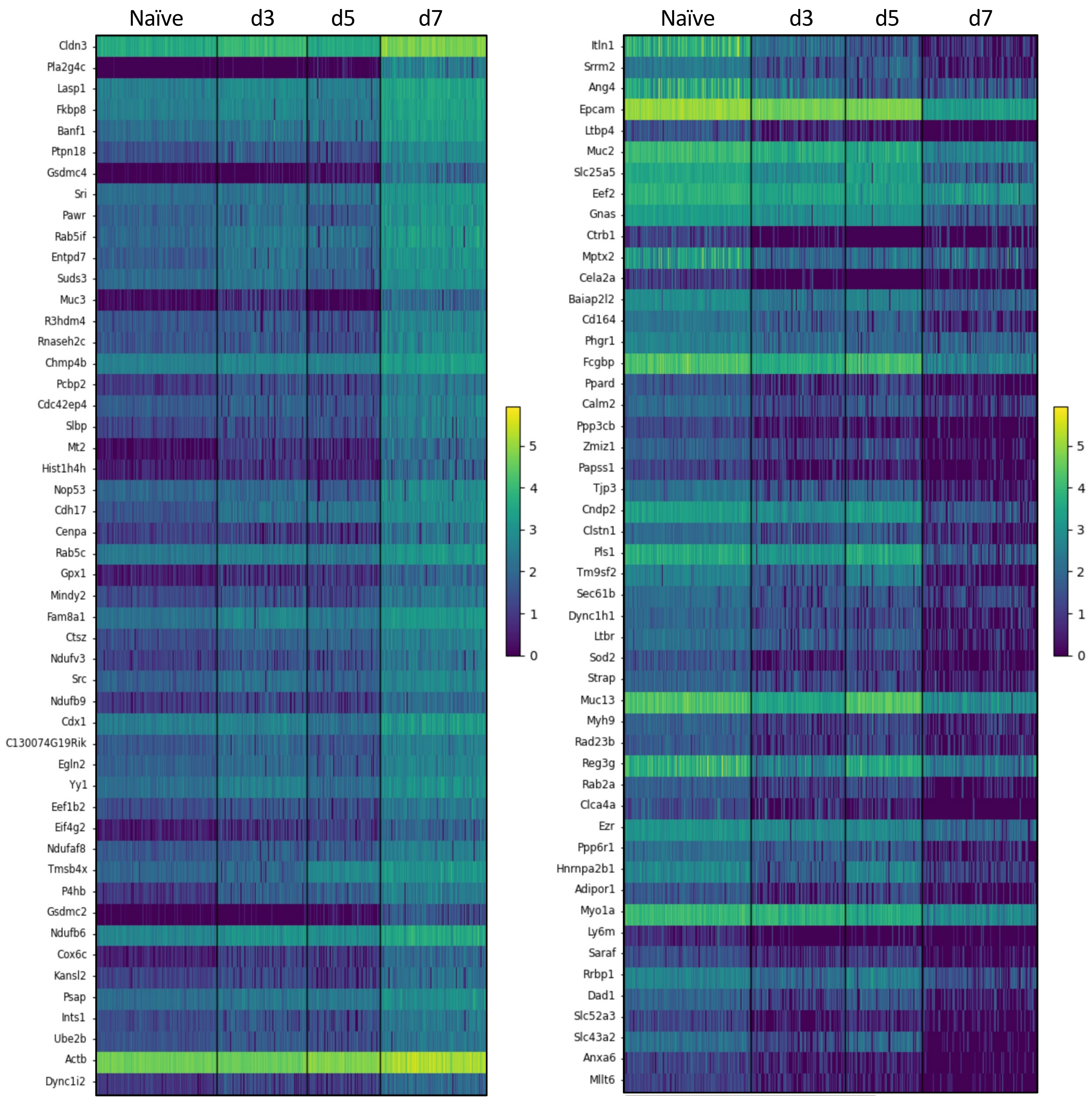

Supplementary Figure 3

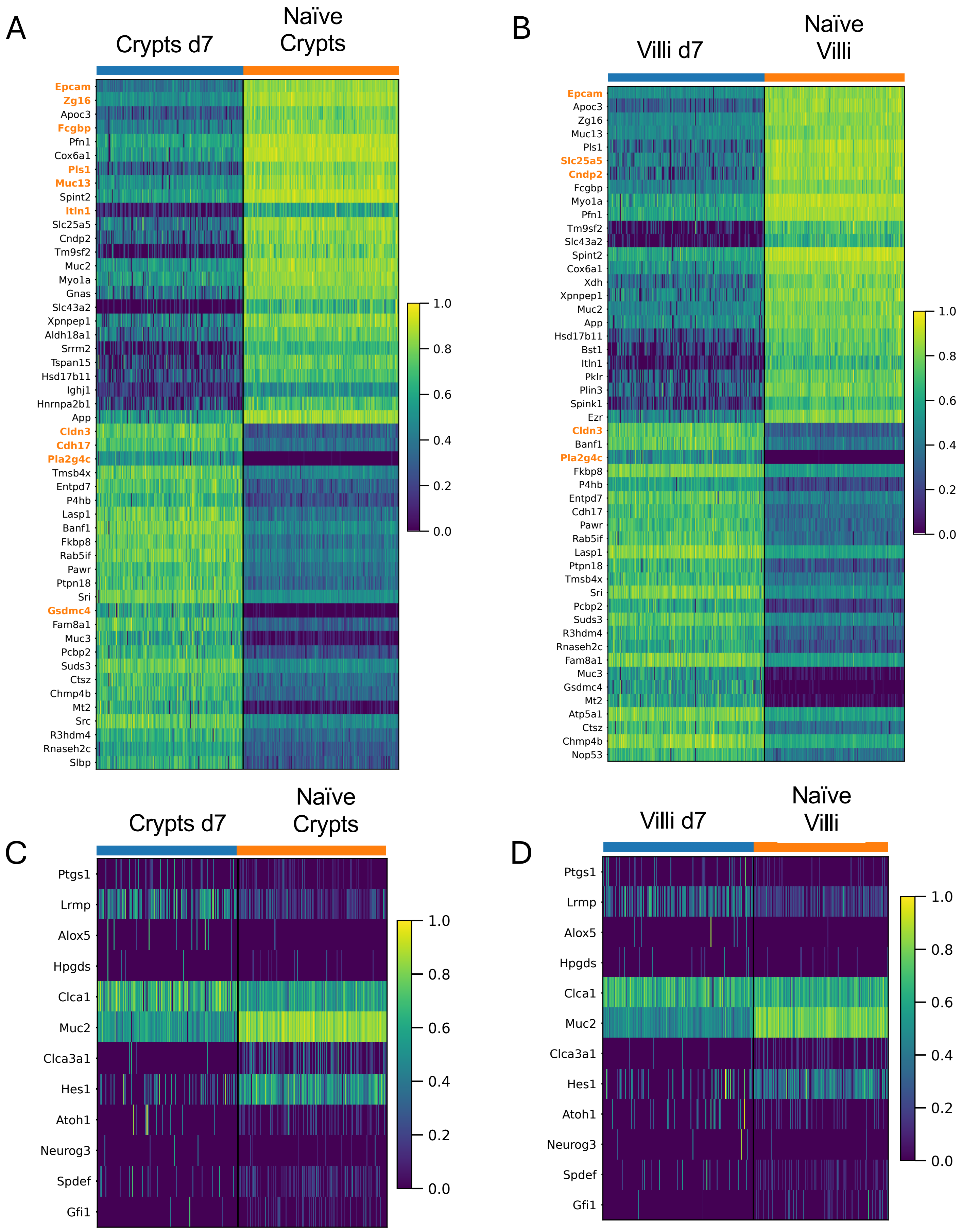

### Supplementary Figure 4

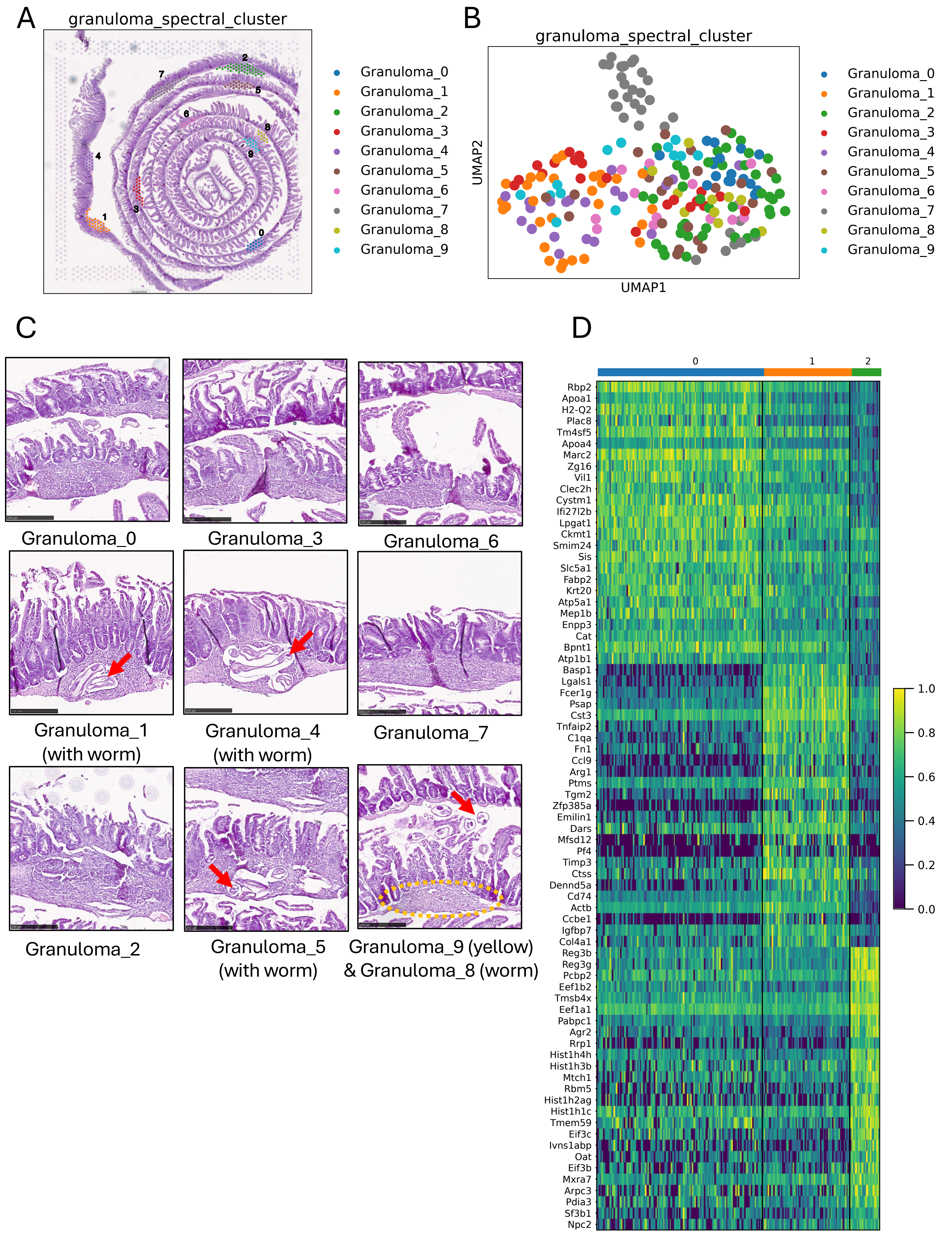

Supplementary Figure 5

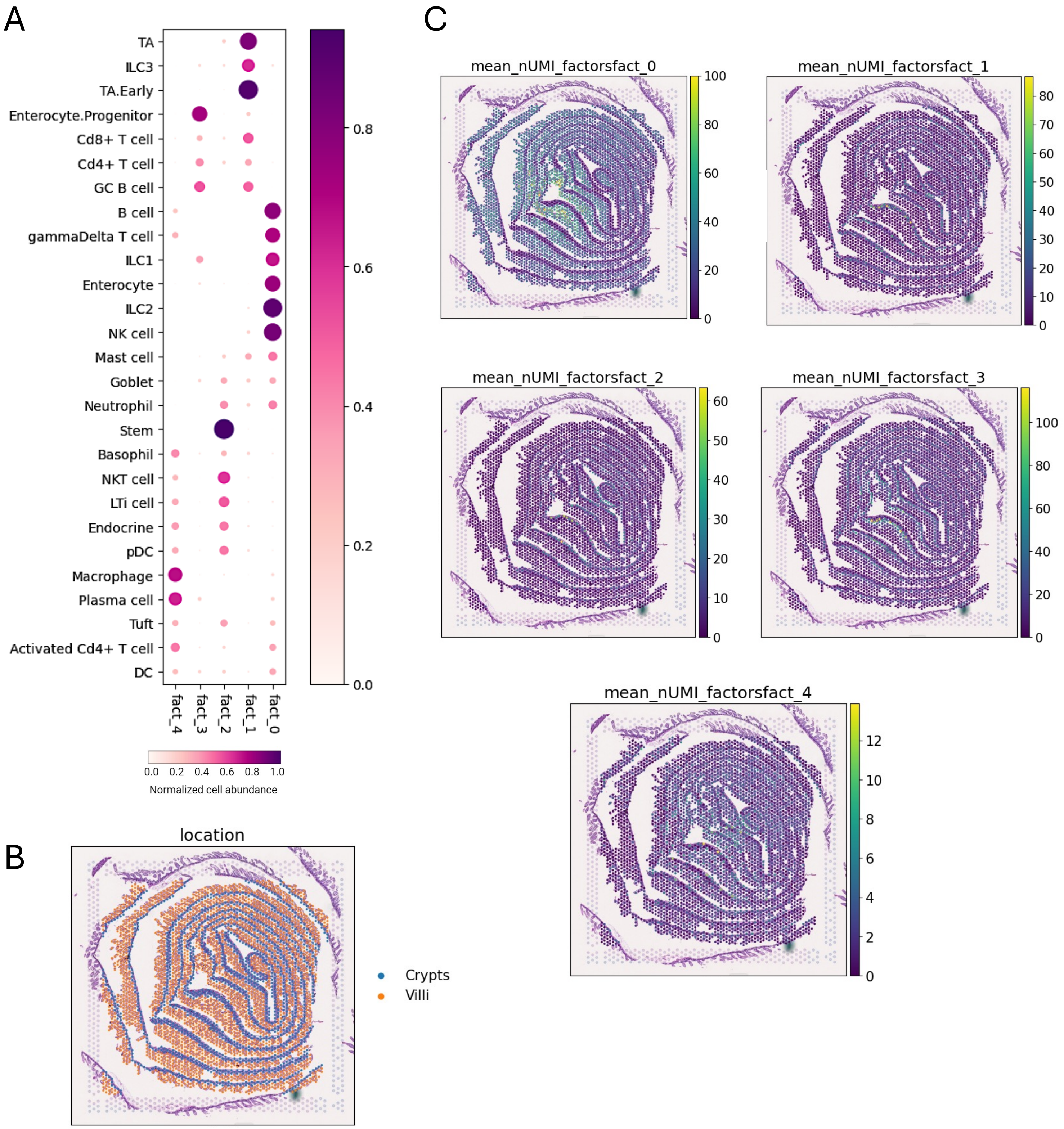

**Supplementary Table 1**

| Antibody | Dilution | Cat. number |
|---|---|---|
| Rabbit anti-RELMα | 1:100 | 500-p214 |
| Rat anti-F4/80 | 1:100 | Ab6640 |
| AF594 anti-rabbit | 1:1000 | A11012 |
| AF647 anti-rat | 1:1000 | A21247 |
| DAPI | 1:1000 | D9542 |
